# Durable Antimicrobial Microstructure Surface (DAMS) Enabled by 3D-Printing and ZnO Nanoflowers

**DOI:** 10.1101/2024.06.11.598554

**Authors:** FNU Yuqing, Shuhuan Zhang, Ruonan Peng, Justin Silva, Olivia Ernst, Blanca H Lapizco-Encinas, Rui Liu, Ke Du

**Affiliations:** Department of Chemical and Environmental Engineering, University of California, Riverside, CA, USA; Department of Mechanical Engineering, University of California, Riverside, CA, USA; Department of Mechanical Engineering, Rochester Institute of Technology, Rochester, NY, USA; Department of Biomedical Engineering, Rochester Institute of Technology, Rochester, NY, USA

## Abstract

Numerous studies have been trying to create nanomaterials based antimicrobial surfaces to combat the growing bacterial infection problems. Mechanical durability has become one of the major challenges to applying those surfaces in real life. In this study, we demonstrate the Durable Antimicrobial Microstructures Surface (DAMS) consisting of DLP 3D printed microstructures and zinc oxide (ZnO) nanoflowers. The microstructures serve as a protection armor for the nanoflowers during abrasion. The antimicrobial ability was tested by immersing in 2E8 CFU/mL *Escherichia coli* (*E. coli*) suspension and then evaluated using electron microscopy. Compared to the bare control, our results show that the DAMS reduces bacterial coverage by more than 90% after 12 hrs of incubation and approximately 50% after 48 hrs of incubation before abrasion. Importantly, bacterial coverage is reduced by approximately 50% after 2 min of abrasion with a tribometer, and DAMS remains effective even after 6 min of abrasion. These findings highlight the potential of DAMS as an affordable, scalable, and durable antimicrobial surface for various biomedical applications.

## B. Introduction

Biofilm is a group of microorganisms that adhere to a surface and produce an extracellular polysaccharide matrix (EPS), and the EPS matrix could increase bacteria’s antibiotic resistance significantly [1]. Biofilms can pose serious threats to human health as they can lead to infections such are sepsis, dental plaque, otitis, cystic fibrosis, endocarditis, and urinary tract [2]. According to the National Institution of Health, biofilms make up about 80% of microbial infections in humans [3]. In the U.S. alone, around 10% of implant surgery patients were affected by biofilm-related infection, which results in around 100,000 deaths annually [4]. Biofilm is not only a problem in the healthcare industry but also in marine industries [5], food industries [6], and more. Developing a robust antimicrobial surface could significantly reduce bacterial infection, especially the concerns of drug-resistance issues [7,8].

Antimicrobial surfaces can be made by applying a coating that limits the growth of microorganisms through chemical processes, modifying the physical structure of the surface to create an environment that is hostile to microbes, and the combination of the two [9]. Ivanova *et al*. demonstrated that black silicon had an effective antimicrobial effect against gram-negative and gram-positive bacteria by inducing stress to the cell membrane to kill the bacteria [10]. Many metal-based antimicrobials have been developed and some have been used in industry applications with gold [11], silver [12], copper [13], and other materials [14]. Although they have shown promising antimicrobial results, the toxicity of metal materials to humans and the environment has raised public attention [14]. TiO_2_ is one of the popular materials for antimicrobial applications. It has been shown that the combination of silver and TiO_2_ nanoparticles had a synergetic effect on its antimicrobial ability with photocatalytic under UV irradiation [15]. However, this mechanism limits its applications such as implants as UV light has a small penetration length. On the other hand, nanomaterials such as Zinc Oxide (ZnO) have been proven to have effective antimicrobial ability for both gram-positive [16,17] and gram-negative bacteria [18,19]. ZnO nanoparticles can be synthesized into many different morphologies using rapid and inexpensive methods [19–22]. ZnO nanostructures release reactive oxygen species (ROS) that can damage the cell wall and efficiently eliminate bacteria [23]. In addition, biomimetic hierarchical nanostructures have been created to reduce bactericidal activity by increasing the liquid-repellent capability at the interface [24].

While nanostructures have demonstrated impressive antimicrobial properties, their limited mechanical durability restricts their applicability in various scenarios. Recently, micropatterns have been developed to protect the anti-wetting materials as robust superhydrophobic surfaces, avoiding damage to the nanostructures by sacrificing the larger size of the micro-frame [25]. In this work, we explore the feasibility of using 3D-printed micropatterns to protect the ZnO nanoflowers as a novel and mechanically robust antimicrobial surface, named Durable Antimicrobial Microstructure Surface (DAMS). We show that DAMS reduces the bacteria on the surface by more than 90% after 12 hrs of incubation. Significantly, DAMS exhibits remarkable antimicrobial efficacy post-abrasion testing, maintaining a 50% reduction in bacteria coverage after 2 min of abrasion and a 20% reduction after 6 min. Since both 3D printing and ZnO growth methods are cost-effective and capable of creating arbitrary shapes in 3D space, our research lays the groundwork for producing a wide array of devices endowed with resilient antimicrobial properties.

## C. Experiments

### Microstructure fabrication

The microstructure chips were designed using SolidWorks. Both inverted pyramids and inverted diamond-shaped structures were created with a feature size of 500 μm and 1,000 μm, respectively [25]. The 500 μm inverted pyramid has a width of 500 μm, while the 1,000 μm inverted pyramid has a width of 1,000 μm. The 500 μm inverted diamond shape has dimensions of 500 μm in length and 300 μm in width, whereas the 1,000 μm inverted diamond shape measures 1,000 μm in length and 600 μm in width. All of them have a depth of 500 μm and the space between each feature (ridge) is 50 μm. The designs were printed using a consumer-graded LCD resin 3D printer (Elegoo Saturn), with “Standard Resin” also from Elegoo. After printing, the microstructure chips were post-processed by placing them into 99% isopropyl alcohol and cleaned with the Elegoo Mercury Plus curing station. Finally, it was cured with 405 nm UV light for 2 min.

### Zinc oxide synthesis and coating

The hydrothermal deposit method was used to synthesize the ZnO nanoflowers. The recipe used in this study contains 0.407 g of zinc nitrate, 1 ml of ammonium hydroxide solution, and 2.5 mL of ethanolamine [26]. These materials were mixed in a beaker with 43 mL of deionized water, and the 3D-printed chips were added to the bottom after. Then, the beaker was placed on a hot plate and heated to 70 °C for 1 hr with mild agitation to allow the ZnO nanoflowers to grow on the 3D-printed chips. The chips were taken out of the solution and airdried overnight before further experiments.

### Bacteria preparation and quantification

*Escherichia coli* (*E. coli*) was obtained from ATCC (25922). A concentration of 2E8 CFU/mL *E. coli* liquid culture media was prepared with brain heart infusion broth. It was quantified by measuring the optical density at 600 nm wavelength using a spectrophotometer (Thermo Scientific™ SPECTRONIC™ 200) and then incubated at 37 °C for 12 and 24 hrs.

### Scanning Electron Microscope (SEM) imaging

SEM was used to quantify the bacteria on the 3D-printed chip. The samples were placed into the Phosphate Buffer Saline solution for 1 hr after the bacteria incubation. Then, they were placed into ethanol for 10 min and then air-dried for 24 hrs. A gold layer was sputter coated onto the sample’s surface for 2 min to reduce the charging effect. SEM images of 83 μm × 83 μm view were taken and then evaluated using ImageJ.

### Abrasion test

The abrasion test was conducted using a self-developed “ring-on-disc” tribometer. Sandpapers (240 grit) were cut into disc shapes and attached to a tribometer. The initial contact force of the tribometer contacting the chip samples was set to 2.5 lbs. Then, the tribometer was rotating at a constant speed of 8 rotations per min. In this study, samples were tested for 2, 4, and 6 min. To ensure the consistency of the abrasion, 240 grit sandpapers were replaced for each test. After the abrasion test, the sample depth of the inverted structures was evaluated using the microscope with a dial test indicator. The focus plane of the top and bottom of the inverted structure was identified and measured with the test indicator. The setup for the abrasion test is shown in Supplementary 1.

## D. Results and Discussion

**Figure 1a** illustrates the overview of the sample preparation processes for DAMS. The 3D-printed substrate was coated with ZnO nanoflowers and the ridge section was scraped to expose the resin surface using a plastic scraper. Then, the abrasion test was performed on the sample surface. Initially, the effectiveness of the ZnO nanoflowers as an antibacterial material was demonstrated. Glass slides with and without ZnO nanoflower coating were incubated with *E. coli* for 12 hrs and the bacteria coverage was evaluated and compared (**Figure 1b**). The results show that under the same area, the bacteria coverage on the glass slide without ZnO coating is ∼3.5 times more than with ZnO coating, indicating that ZnO nanoflowers can prevent bacteria attachment on the solid surface (**Figure 1c**). **Figure 1d** demonstrates the 3D printed structure’s ability to protect the ZnO nanoflowers. Before scraping and rubbing the surface with a plastic scraper, a significant amount of ZnO is grown in the 3D-printed microstructures as the surface shows a white color. After scraping and rubbing, there are still significant amounts of ZnO nanoflowers remaining on the 3D printed micropatterned structure structures, mostly in the valley of the microstructures.

**Figure 1.**
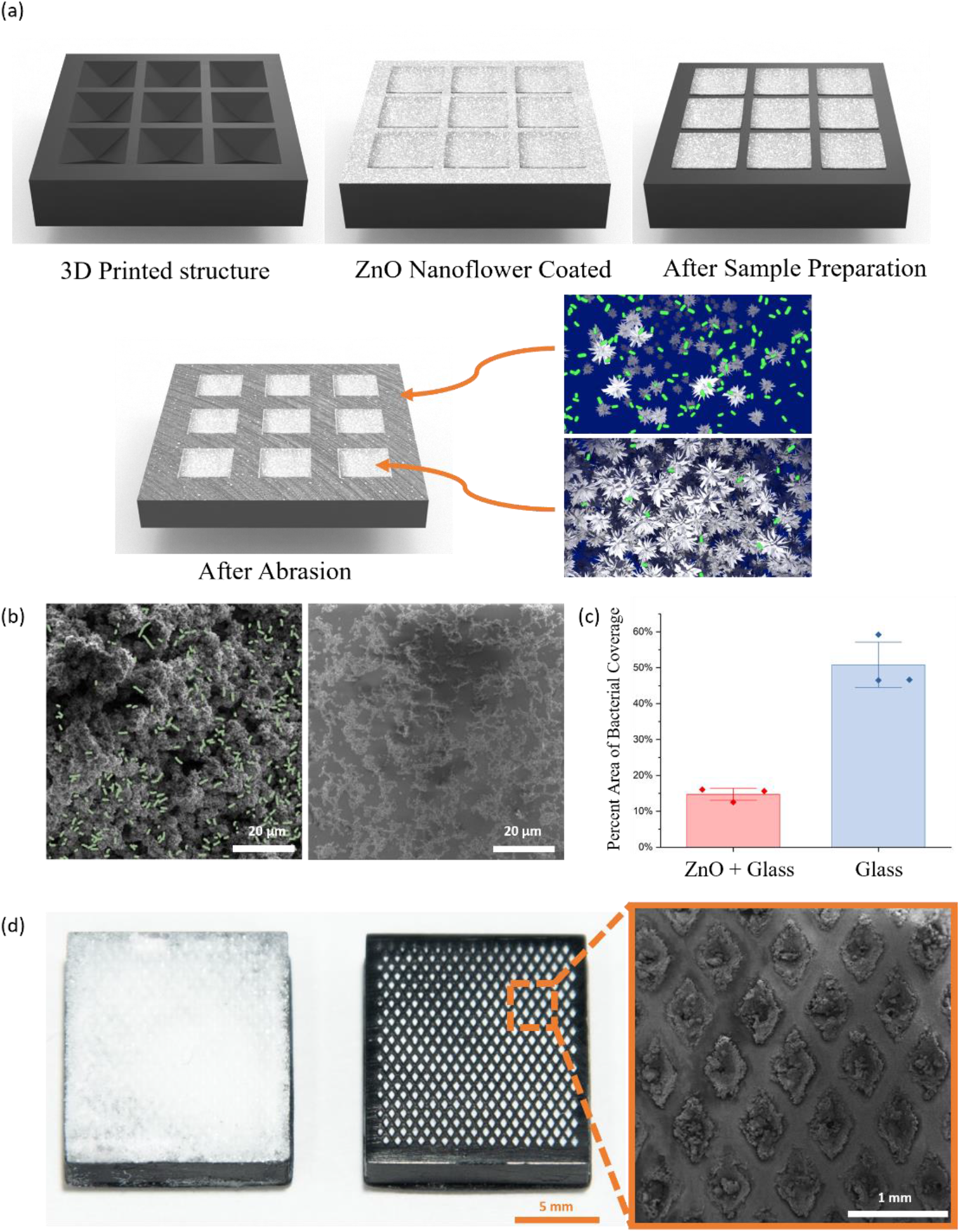
(a) Illustration of the 3D printed structure after coated with ZnO nanoflower, sample preparation, and abrasion. (b) SEM images of the glass side with (left) and without (right) the ZnO coating after 12 hrs of *E. coli* incubation. (c) Percent area of bacteria coverage of glass sides with and without ZnO coating. (d) Micrograph image of the 3D printed microstructures after ZnO deposition (left) and initial treatment by removing the ZnO on the ridge (middle). SEM image of the microstructures after ZnO removal on the ridge (right).

**Figures 2a** and **2b** show the SEM images of the inverted pyramid structures after 12 hrs of *E. coli* incubation with and without the ZnO nanoflower coated on the microstructures, respectively. Without ZnO nanoflowers, it was observed that most of the *E. coli* would concentrate on each valley section of the inverted structure. The ridge section is relatively smooth and thus has less bacteria adhesion. With ZnO nanoflowers, the *E. coli* coverage of the valley section decreased significantly. The percentage area of bacteria coverage results of both square and diamonded shapes with two different sizes at the valley section are presented in **Figure 2c**. The 500 μm feature-sized micropatterned structures were able to reduce up to 90% of bacteria coverage with ZnO nanoflower coating after 12 hrs of incubation compared to the control. It suggests that the smaller feature size, compared between 1,000 μm and 500 μm, would increase the chip’s *E. coli* inhibition ability. The 1,000 μm feature-sized micropattern structure still reduced the bacteria coverage by more than 60%. On the other hand, the difference between the diamond and square-shaped micropatterned 3D printed structures is relatively small, without showing significant discrepancies.

**Figure 2.**
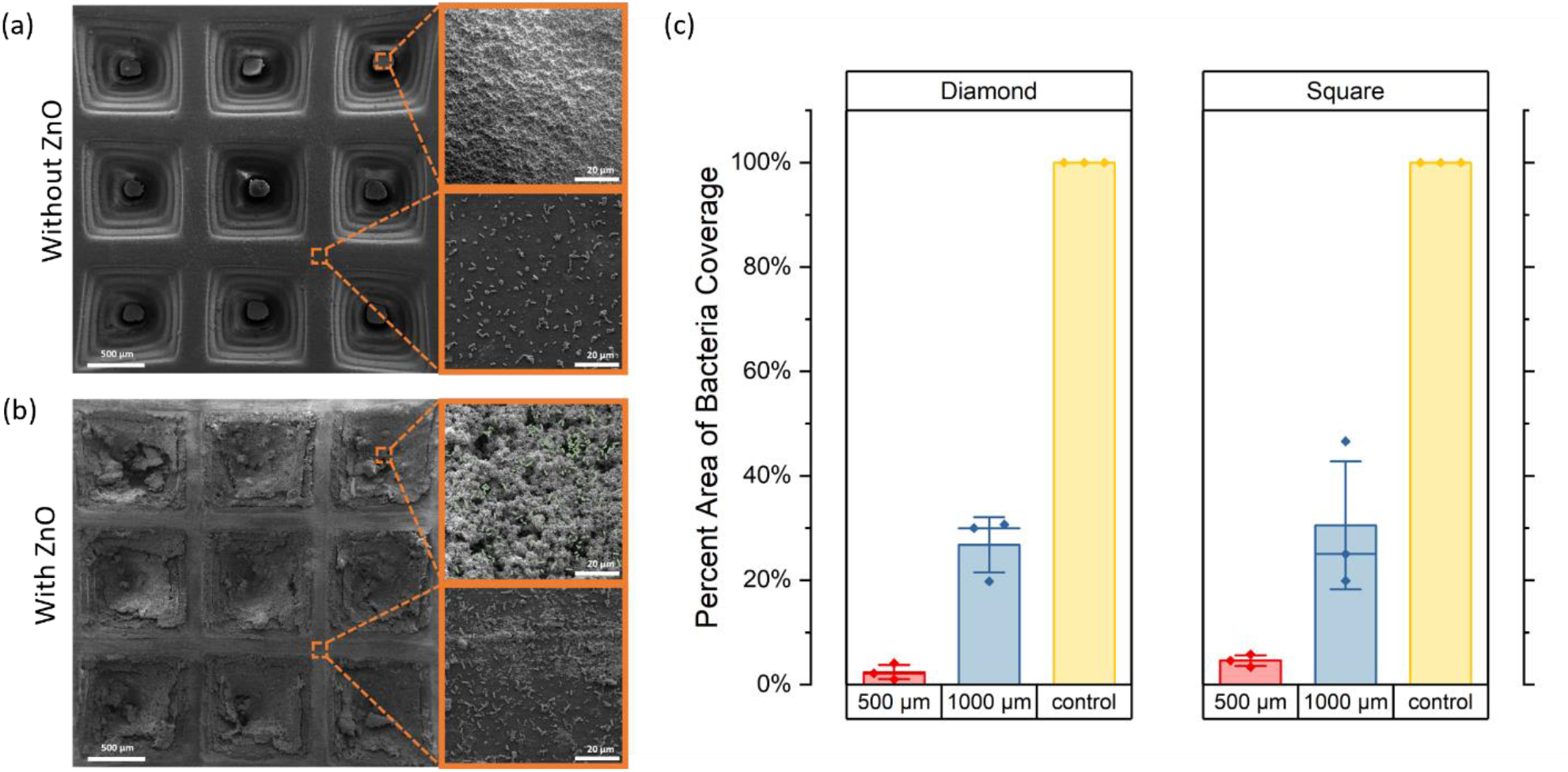
SEM images of the 1,000 μm inverted pyramids structured chip (a) without ZnO coating after 12 hrs of *E. coli* incubation and (b) with ZnO coating after 12 hrs of *E. coli* incubation. (c) Percent area of bacteria coverage on the valley section after 12 hrs of incubation with both diamond and inverted pyramids patterned chips. The control sample is a 1,000 μm diamond patterned chip without ZnO coating.

After demonstrating that the ZnO nanoflower coating and the micropatterned structures are effective in reducing bacteria adhesion, the durability of the DAMS was evaluated through the abrasion test. A tribometer was used to produce abrasions in the samples consistently. The SEM image of the squared micropatterned chip without ZnO coating after 2, 4, and 6 min of abrasions is shown in **Figure 3a**. As expected, the surface of the structure became rougher over time, and the top dimension of the inverted structure became smaller as the abrasion time increased. Due to the tribometer device being circular, the scratch marks on the sample chip appeared to be concentric. The depth of the inverted structures after the abrasion was measured and shown in **Figure 3b**. The depth of the structure was reduced from ∼0.09 to ∼0.002 inches after 6 min of abrasion. **Figure 3c** shows the SEM images of the ridge and valley section of the ZnO-nanoflower-coated samples after 2, 4, and 6 min of abrasion. It was observed that the ZnO nanoflowers initially located inside of the valley got transported to the ridge section of the chips as the abrasion time increased. For the 6-min sample, the amount of ZnO nanoflowers on the ridge and in the valley section are comparable.

**Figure 3.**
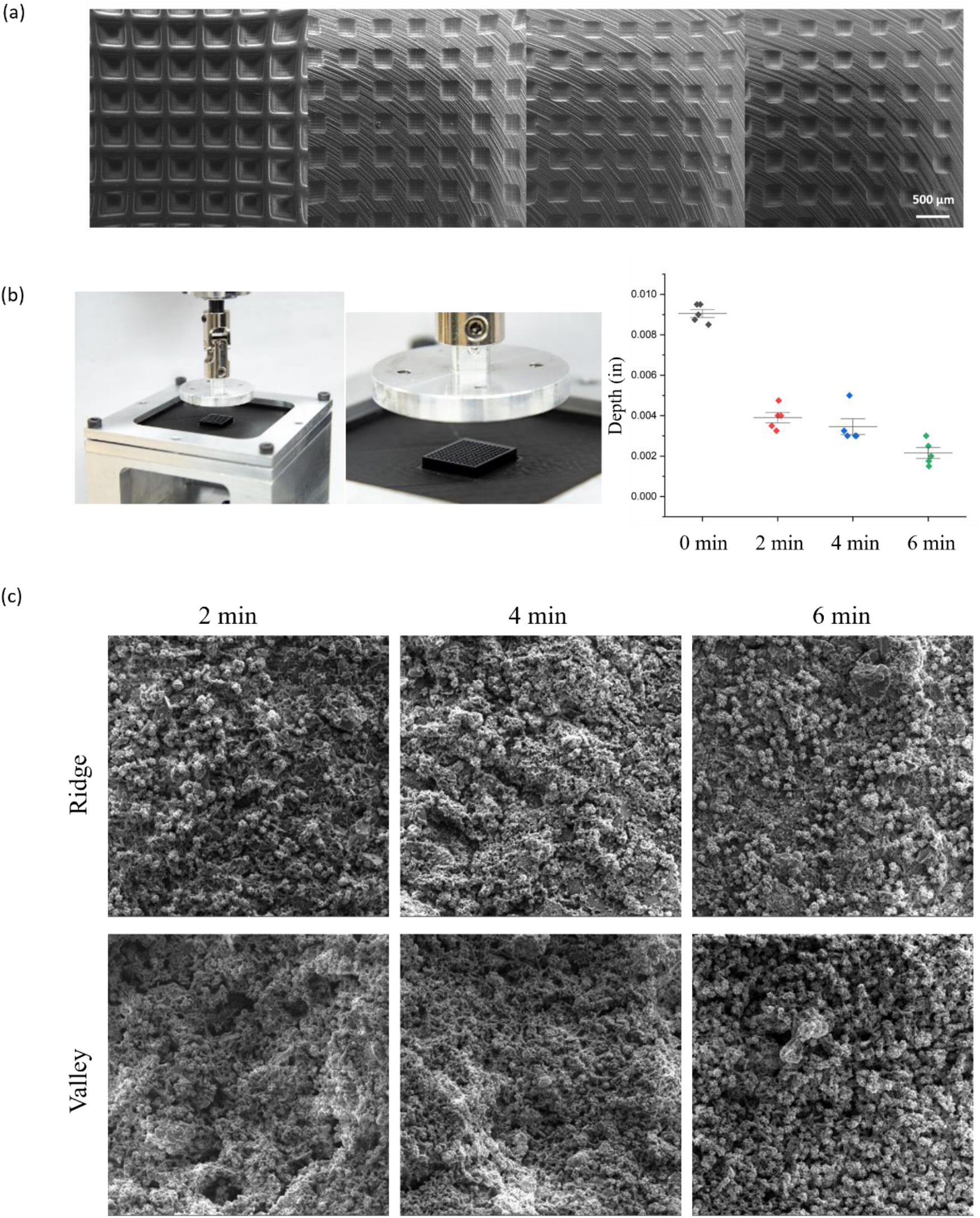
(a) SEM images of the 3D printed microstructures (before ZnO coating) with an abrasion test time ranging from 0 to 6 min. (b) Abrasion setup and depth of the structure after abrasion. (c) SEM images of the ZnO nanoflower coated samples’ ridge and valley area after abrasion and then 12 hrs of *E. coli* incubation.

We tested the *E. coli* coverage on the abrased samples after 12 hrs of incubation. After 2 min of abrasion, the DAMS achieved 80% bacteria reduction, showing the robustness of our surface. As predicted, the number of bacteria cells increased in the valley sections with increasing abrasion time. After 6 min of abrasion, the bacteria coverage results were still about 20% better than the control which has no ZnO nanoflower coating nor abrasion, as shown in **Figure 4a**. The ridge section, on the other hand, shows that the number of bacteria cells decreases after the abrasion compared to the samples before the abrasion (**Figure 4b**). This matches our observation that the ZnO nanoflower is being transported onto the ridge during abrasion, as shown in **Figure 4c**. This phenomenon increases the overall antimicrobial performance of our surface as ZnO nanoflowers can better reduce bacteria growth than the smooth resin surface. It also explains why the control has fewer bacteria cells on the ridge compared to the one without abrasion (0 min). **Figure 5** shows the surface with a 500 μm feature size of both diamond and square-shaped structures’ percent area coverage of bacteria after 48 hrs of *E. coli* incubation. After 2 days of incubation, DAMS continues to reduce the percent area coverage of bacteria by approximately 50%, demonstrating that our chip can effectively slow down biofilm formation over an extended period.

**Figure 4.**
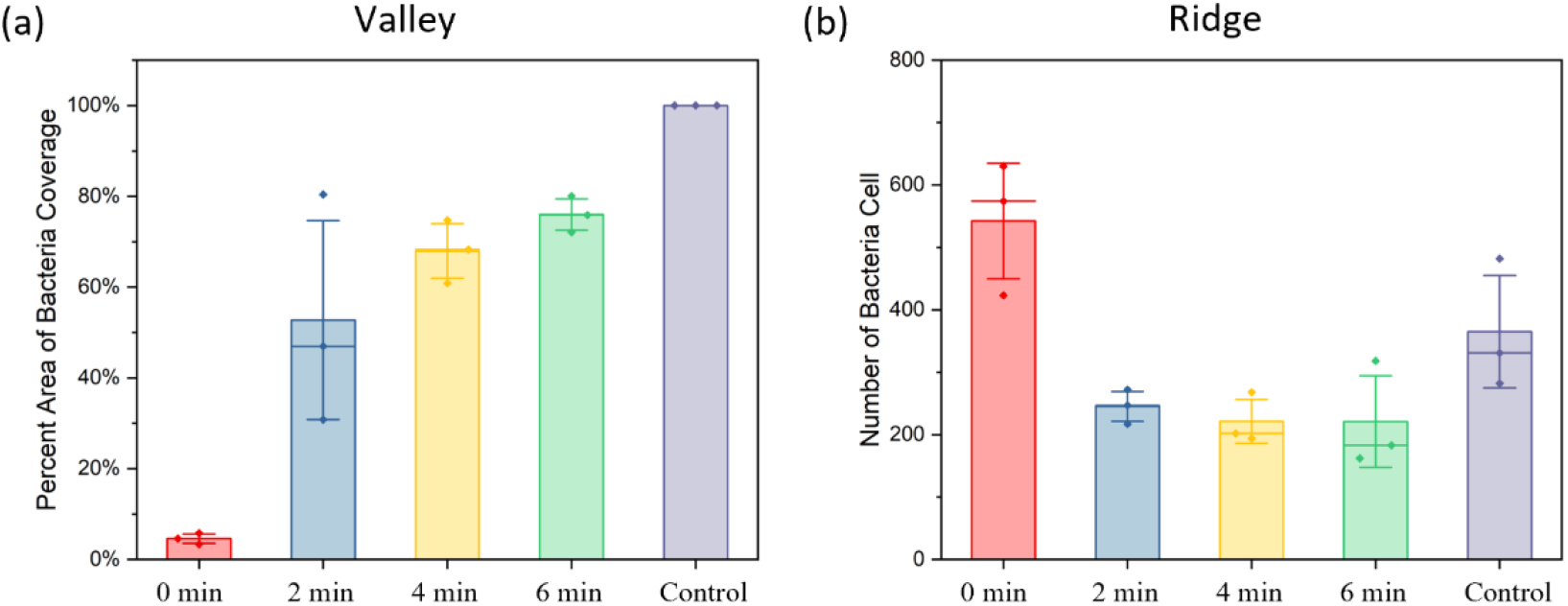
Results of (a) valley and (b) ridge section of the ZnO nanoflower coated square samples (feature size: 500 μm) incubated with *E. coli* for 12 hrs after abrasion tests from 0 to 6 min.

**Figure 5.**
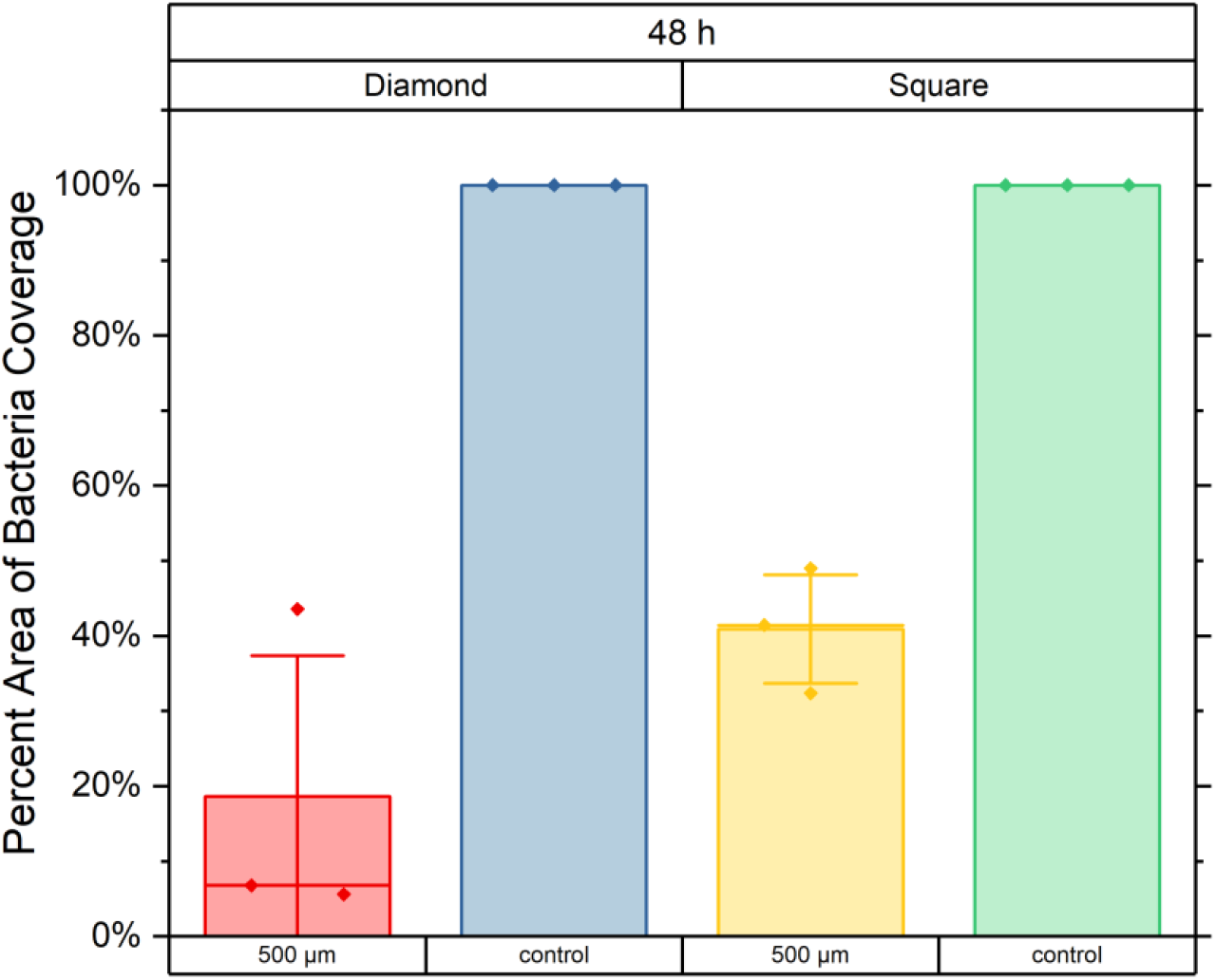
Percent area of bacteria coverage of diamond and square-shaped structure with 500 μm feature size after 48 hrs of *E. coli* incubation.

## E. Discussion

The abrasion resistance was reported to be one of the biggest obstacles for the nanoscale-based antimicrobial surfaces developed in laboratories to transform into clinical applications [6,26]. We developed a durable antimicrobial surface with the combination of the micropatterned 3D printed structure with ZnO nanoflower coating. We demonstrated an inexpensive and easy way of fabricating the surface. Only consumer-graded equipment of less than $500 was needed to obtain the structure and coating shown in this study, instead of expensive clean room equipment used in some studies [10,27]. The DAMS, measuring 13 mm by 15 mm by 3 mm, costs less than $0.02, making it easily scalable for various applications. Compared to other micro or nano-scaled antimicrobial surfaces, our developed surface demonstrates excellent wear resistance [10,27].

Even after 6 min of tribometer abrasion, it still outperforms the uncoated control in bacterial inhibition.

One significant challenge in creating a durable and effective antimicrobial surface lies in preventing material removal during abrasion. Re-depositing antimicrobial materials after use is often necessary but impractical in numerous settings. The structures developed such as black Si [10] or TiO_2_ micropillar [27] would get damaged after abrasion and lose their antimicrobial ability, and surface treatment such as superhydrophobic coating has no exception [28]. In our case, we found that the antimicrobial agent, ZnO nanoflowers, originally stored in the valley section of the structure would get transported on the surface, as shown in **Figure 3C**. This effect would prolong the antimicrobial surface’s lifetime even after it’s been abrased making it a more economical choice in long-term applications.

The antimicrobial surface proposed in this study has a flexible manufacturing process and is not limited to one type of nanomaterial coating. The structure could be injection molded or embossed for mass production using plastic material [29]. Metal material could also be selected for the microstructure [30]. Depending on the type of metal and coating, the combination could have synergetic effects on its antimicrobial ability. Many studies have demonstrated that the ZnO nanostructures could be combined with resin or coating to further improve their durability [31– 33]. Therefore, DAMS not only works with ZnO nanoflowers, but it has the potential to be coated with other materials depending on the different applications such as titanium dioxide nanoparticles and gold nanoparticles [34].

This study only demonstrates robust antimicrobial properties on a flat surface. The emerging high-resolution 3D printing techniques can fabricate high-resolution microstructures in 3D space and ZnO can be grown on every surface in the solution which is uniform and scalable [35,36].

Therefore, in the future, the inverted surface could be made into curved surfaces which would enable more application areas such as braces in the dental industry and urinary catheters [37,38]. In addition, ZnO was reported to have little to no toxicity and is biocompatible [39–41] which is ideal for long-term bacterial inhibition.

## F. Acknowledgments

This work is partially supported by the National Institutes of Health (Grant No. R35GM 142763).

