## Supplement for "Durable Antimicrobial Microstructure Surface (DAMS) Enabled by 3D-Printing and ZnO Nanoflowers"

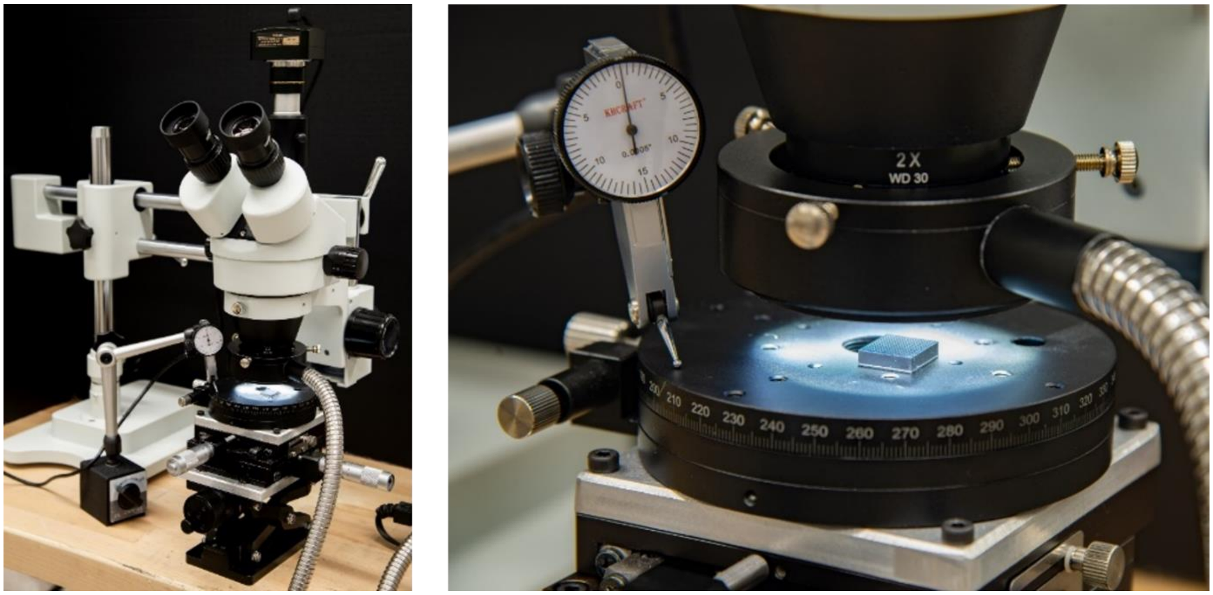


**Supplementary 1.** Depth measurement set up for the DAMS.
